# Before the eye moves: microsaccade preparation expands spatial integration at the center of gaze

**DOI:** 10.64898/2026.07.09.737553

**Authors:** Krishnamachari S. Prahalad, Martina Poletti

**Affiliations:** Department of Brain and Cognitive Sciences, University of Rochester; Center for Visual Science, University of Rochester; Department of Neuroscience, University of Rochester

## Abstract

Fixation is often treated as a period of stable visual processing. Yet, fixation is often punctuated by frequent microsaccades that occur during tasks involving complex foveal stimuli. These small eye movements are preceded by changes in visual sensitivity, both at the upcoming movement goal and at the currently fixated location. However, previous work has largely focused on isolated stimuli, leaving unclear whether pre-microsaccadic modulations reflect changes in sensitivity alone or also alter the spatial interactions that govern object recognition. Visual crowding provides a direct test of this question because it depends on the integration and segregation of nearby features and constrains recognition even within the foveola. Using high-precision Dual Purkinje Image eye tracking with retinally contingent stimulus delivery, we measured acuity and crowding thresholds at the preferred locus of fixation (PLF), the starting point of the impending gaze shift, while observers either maintained fixation or prepared to execute a microsaccade to a cued location. Unflanked acuity at the PLF remained stable across conditions. In contrast, crowding strength increased during the pre-microsaccadic interval, indicating an expansion of the foveal crowding zone. These results show that microsaccade preparation alters spatial integration at the starting point of the movement, increasing crowding even when sensitivity to isolated stimuli remains unchanged. Thus, microsaccades reshape foveal vision not only by modulating visual discrimination at the movement goal, but also by changing how nearby features are integrated and segregated before the eyes move.

**Significance Statement:** Vision is often assumed to be most stable when gaze is fixed. Yet the eyes are never truly still, and the brain continually prepares small movements that shape perception before they occur. This study shows that such preparation changes how visual information is organized at the very center of gaze. Upcoming eye movements do not simply alter sensitivity to isolated objects; instead, they change how nearby features are integrated. These findings reveal that fine spatial vision and object recognition depends not only on what falls on the retina or on subsequent cortical processing but also on what the eyes are preparing to do next.

## Introduction

Foveal vision during fixation is often treated as a period of stable visual processing, but fixation is punctuated by frequent microsaccades. These small eye movements occur during a wide range of high-acuity tasks, including reading, face recognition, optotype identification, and fine visuomotor coordination (Bowers and Poletti, 2017, Ko et al., 2010, Shelchkova et al., 2019, Intoy and Rucci, 2020, Clark et al., 2022). Microsaccades are closely linked to visual analysis: they reposition fine spatial detail on the retina and are preceded by changes in visual sensitivity and attention (Rucci and Poletti, 2015, Guzhang et al., 2024, Shelchkova and Poletti, 2020, Prahalad and Coates, 2024). Understanding how these movements modulate perception is therefore essential for a complete account of fine spatial vision.

Recent work has shown that, before microsaccade onset, visual processing is modulated both at the upcoming movement goal and at the preferred locus of fixation (PLF), the starting point of the impending gaze shift. At the movement goal, processing resources are allocated with high spatial precision, enhancing discrimination within a restricted region around the microsaccadic target (Guzhang et al., 2024). At the PLF, by contrast, performance for isolated stimuli can decline during the pre-microsaccadic interval (Guzhang et al., 2024), consistent with a shift in processing priority away from the currently fixated location. Similar reductions in foveal sensitivity have been reported before larger saccades (Hanning and Deubel, 2023, Kroell and Rolfs, 2022). However, prior work has largely characterized these modulations using isolated stimuli. It therefore remains unclear whether pre-microsaccadic changes at the PLF reflect only a change in visual sensitivity, or whether they also alter the spatial interactions that govern object recognition.

Visual crowding provides a direct way to address this question. Crowding impairs recognition when nearby features fall close to a target, even without physical overlap (Stuart and Burian, 1962, Bouma, 1970). It reflects the balance between integration and segregation of adjacent visual information (Levi, 2008, Pelli, 2008, Whitney and Levi, 2011), and constrains recognition even within the foveola, where visual acuity is highest (Flom et al., 1963, Coates and Levi, 2014, Siderov et al., 2013, Bondarko et al., 2024, Clark et al., 2024, Shamsi et al., 2022, Prahalad et al., 2025). Notably, in natural scenes, this same integration facilitates the perception of textures, contours, and global patterns through Gestalt-like grouping mechanisms (Manassi et al., 2012, 2013, Herzog et al., 2015). Thus, crowding is not merely a nuisance but a consequence of a visual system optimized to balance the integration and segregation of information. Even within the foveola, where visual acuity is highest, crowding imposes a fundamental limit to object recognition (Pelli et al., 2016, Flom et al., 1963, Coates and Levi, 2014, Siderov et al., 2013, Bondarko et al., 2024, Clark et al., 2024, Shamsi et al., 2022, Prahalad et al., 2025). Thus, assessing crowding during microsaccade preparation makes it possible to distinguish changes in sensitivity to isolated stimuli from changes in the spatial integration of nearby features.

For larger saccades, crowding has been reported to diminish at the saccade goal before movement onset (Harrison et al., 2013, Lin et al., 2014, Wolfe and Whitney, 2014), yet this effect remains debated because subsequent work suggested a more general enhancement of performance rather than a crowding-specific modulation (Ăgaŏglu et al., 2016, Ăgaŏglu and Chung, 2017, Buonocore et al., 2017). Within the foveola, pre-microsaccadic reduction of visual crowding has also been observed at the future microsaccadic goal, with some evidence that the geometry of the crowding zone may be altered before the eye moves (Prahalad and Coates, 2024). What remains unknown is what happens at the PLF, where the microsaccade begins. This question is important because increased crowding at the PLF would indicate that microsaccade preparation reshapes spatial integration at the currently fixated location, rather than merely changing visual sensitivity.

Crowding provides leverage beyond unflanked acuity because it reveals how nearby features are selected, suppressed, or pooled. A pre-microsaccadic reduction in performance for flanked targets at the PLF could arise from several sources: increased internal noise, which should impair both unflanked and flanked acuity; reduced suppression of flankers, which should increase target–flanker confusions; or enlarged spatial pooling regions, which should selectively increase crowding strength and critical spacing even when unflanked acuity remains stable (Lu et al., 2011, Pelli, 1990, Greenwood et al., 2009, Parkes et al., 2001, Pelli et al., 2004, Levi, 2008). To distinguish these possibilities, we measured both unflanked acuity and flanked thresholds at the PLF while observers either maintained fixation or prepared and executed a microsaccade. We used high-precision Dual Purkinje Image eye tracking (Wu et al., 2023) with gaze-contingent stimulus delivery (Santini et al., 2007), allowing stimuli to be presented with arcminute precision relative to the PLF. This approach enabled us to test whether pre-microsaccadic modulations at the start site reflect a general change in visual sensitivity or a more specific alteration of foveolar crowding and spatial integration.

## Results

To assess if and how microsaccade planning and execution impact the magnitude of visual crowding at the preferred locus of fixation, we measured visual acuity thresholds in the presence and absence of flankers either during maintained fixation (baseline condition) or immediately before the onset of instructed microsaccades (pre-microsaccade condition). Visual acuity was measured using a 4-alternative forced-choice (4AFC) digit discrimination task. Stimuli consisted of digits in Pelli font, specifically designed to evaluate foveal crowding (Pelli et al., 2016) (see figure 1*A*). In the flanked condition, flanker digits were positioned along the horizontal axis, with center-to-center spacing set to 1.4 times the target size, ensuring that spacing scaled proportionally with stimulus size.

**Figure 1:**
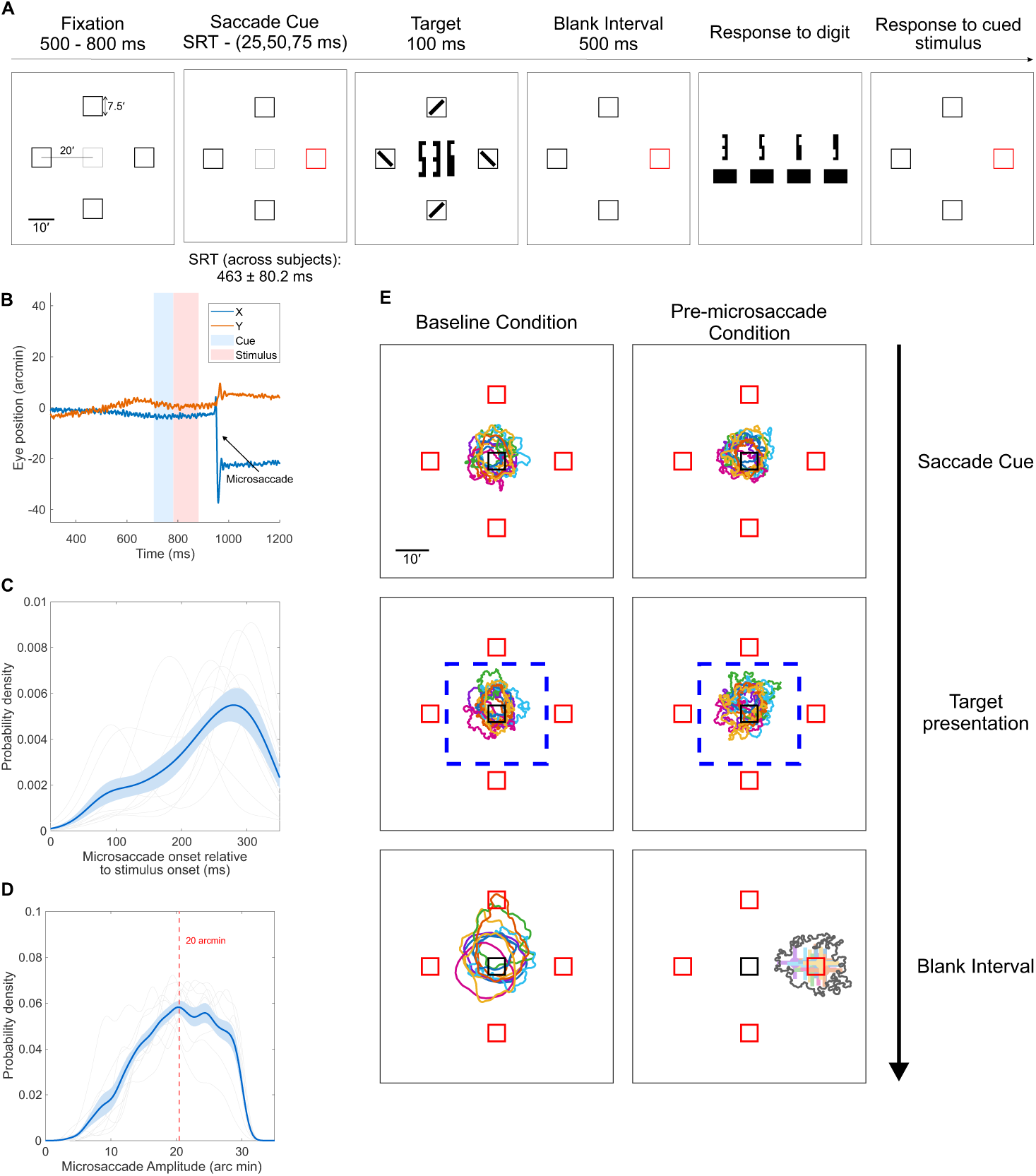
Experiment protocol and characterization of gaze behavior. (A) Experimental protocol illustrating the trial sequence. (B) Example eye trace from the pre-microsaccade condition, showing horizontal and vertical gaze position (arcmin) as a function of time, with the stimulus presented prior to microsaccade onset. (C) Distribution of microsaccade onset times relative to stimulus onset for valid pre-microsaccade trials. (D) Distribution of microsaccade amplitudes. The dashed red line marks the median amplitude (20 arcmin). The blue line and shaded region indicate the across-subject mean *±* SEM, and the shaded black lines show individual subject data. (E) Gaze behavior for baseline (left column) and pre-microsaccade trials (right column) across different time windows in the experimental sequence, represented as 2D gaze contour maps encompassing 68% of gaze positions. Each row corresponds to a specific trial epoch: the saccade cue interval, the target presentation interval, and the subsequent blank interval. The blue dashed square during target presentation marks the 15 *×* 15 arcmin gaze selection window used for trial inclusion. During the blank interval, subjects either maintained fixation close to the center of the display (baseline condition) or executed a microsaccade toward the cued direction (microsaccade condition). The bottom-right panel shows microsaccade landing positions measured 10–30 ms following microsaccade offset. Colored markers denote each subject’s mean landing position, with error bars indicating *±* 1 standard deviation across trials. The overlaid gray contour represents the two-dimensional distribution encompassing 68% of landing positions pooled across subjects. To aid visualization, landing positions were rotated such that all data are shown relative to a rightward cued location.

As microsaccades are typically executed to re-center gaze toward behaviorally relevant details in complex foveal visual input, fine spatial stimuli were presented at four peripheral locations (20 arcminutes from the center of the display) to engage microsaccade planning. Gaze position was monitored using a high-precision eye tracker coupled with a custom gaze-contingent display (Wu et al., 2023, Santini et al., 2007), enabling accurate localization of the line of sight and online detection of microsaccadic activity (Poletti and Rucci, 2016) (see Figures 1*B*, *C* and *D*). A peripheral motor cue—one of four square placeholders changing color from black to red—signaled the target location and instructed observers to prepare a microsaccade; cue–stimulus timing was adjusted based on individual microsaccadic latency estimates to ensure that stimuli were presented prior to microsaccade onset within a 0–350 ms window (see methods for details). In the baseline condition, subjects maintained central fixation throughout the trial and covertly attended to the cued location, as indicated by the change in the placeholder squares’ color. Stimuli were briefly presented and consisted of a target digit surrounded by two flankers at the preferred locus of fixation together with a high-acuity oriented line at each of the off-center locations in the array; importantly, this stimulus disappeared before eye movement execution and did not remain on the display after microsaccade landing. Observers were required to first report the identity of the central digit and then the orientation of the tilted line at the cued location on each trial, ensuring engagement at the microsaccade target location. Thus, the motor cue served both to elicit microsaccade preparation and to define the task-relevant location with 100% cue validity. Analyses were restricted to trials in which retinal input at the time of target presentation was comparable across conditions. To this end we only retained trials in which gaze remained within a central 15 *×* 15 arcmin window during target presentation and trials with no blinks or saccades between cue and stimulus onset. Further, in pre-microsaccade condition, only trials in which a microsaccade occurred within 0–350 ms after stimulus onset, and landed within 15 arcmin from the cued location, were selected. In the baseline condition only trials without microsaccadic events within 300 ms following stimulus onset were selected for analysis (see methods for details). A summary of the number of valid and rejected trials for each observer and condition is provided in Supplementary Table 1.

Figure 1*E* shows the distribution of gaze positions for each subject (separate colors) at different time points (rows) in the trial, separately for baseline and pre-microsaccade conditions (columns). The blue dashed square marks the 15 *×* 15 arcminute gaze window used as the criterion for trial selection. Gaze was maintained close to the center of the display during stimulus presentation in both conditions. The average distance from the display center was highly similar in the baseline (5.64 ± 0.87 arcmin) and pre-microsaccade conditions (5.86 ± 0.80 arcmin). Although gaze was slightly farther from the display center in the pre-microsaccade condition, the magnitude of this difference was small, corresponding to a median increase of only 0.16 arcmin (IQR = 0.24; Wilcoxon signed-rank test: *W* = 6.00, *p* = 0.027). In contrast, fixation stability, quantified by the 68% bivariate contour ellipse area (BCEA), did not differ between conditions (pre-microsaccade: 224.98 ± 68.03 arcmin^2^; baseline: 216.70 ± 63.24 arcmin^2^; *t*(9) = −0.97, *p* = 0.36, Cohen’s *d* = −0.31, *BF*_10_ = 0.46). The bottom row of Figure 1*E* shows gaze position during the blank interval. In the baseline condition, gaze remained centered, whereas in the pre-microsaccade condition, microsaccades consistently landed near the cued location following stimulus offset. This pattern confirms accurate execution of the instructed gaze shifts and indicates that the trial selection and landing criteria effectively isolated valid pre-microsaccade trials.

We first verified that subjects could reliably perform the orientation-discrimination task for the stimuli presented at the microsaccade goal location. To this end, we varied the line’s stroke width between 1 and 1.5 arcminutes - a limited range chosen to prevent the orientation probe from overlapping with the placeholder box. Although this range was constrained by spatial layout considerations, it allowed us to confirm that subjects performed above the 2AFC guess rate (50%) in the baseline condition (mean proportion correct: 0.66 ± 0.07). Most subjects achieved accuracies around 70%. For three participants, performance remained close to chance in the baseline condition even with the largest stimulus setting; however, for two of these participants, performance improved in the pre-microsaccade condition. Only one participant remained near chance across both conditions. Importantly, these results indicate that the large majority of observers were actively engaged with both the central digit and peripheral orientation tasks.

We then compared performance at the cued location between baseline and pre-microsaccade conditions to test whether preparing and executing a microsaccade enhances performance at the upcoming saccade target, reflecting a transient redistribution of processing resources before the gaze shift, as reported previously Guzhang et al. (2024). Figure 2 shows both the proportion correct and the corresponding *d^′^* estimates for the orientation-discrimination task. Consistent with previous work (Shelchkova and Poletti, 2020, Guzhang et al., 2024, Prahalad and Coates, 2024), performance at the cued location was significantly higher in the pre-microsaccade condition compared to the baseline condition (t(9) = −3.56, p = 0.006, Cohen’s d = 0.74, *BF*_10_ = 9.16). On average, performance improved by 6.0% ± 5.7 and *d^′^* increased by 0.35 ± 0.31 in the pre-microsaccade condition relative to baseline. This shows that the performance benefit was specific to trials in which a microsaccade was imminent. Importantly, the enhancement cannot be explained simply by the deployment of covert attention to the cued location, because the cue was 100% valid also in the baseline condition. This result indicates that processing resources were effectively directed toward the microsaccade goal. Once we confirmed that microsaccades were associated with the enhancement of performance at the goal location, we next asked how this preparatory process influences perception at the current fixation site. While most previous studies have focused on perceptual benefits at the microsaccade goal, our interest was in whether this attentional Figure 3*A* shows average psychometric fits across all subjects, plotted separately for the baseline and pre-microsaccade conditions. Individual subject fits are shown in Supplementary Figure S1. Solid lines represent the mean curves, generated by averaging the predicted proportion correct from each subject’s fitted psychometric function (see methods section), and the surrounding shaded areas indicate the standard error of the mean (SEM). We first verified that the current paradigm reliably elicited foveal crowding during fixation. Specifically, we compared visual acuity thresholds between unflanked and flanked trials at the center of gaze (Figure 3*B*, left panel), reported here as target stroke width for the Pelli font (Clark et al., 2022, 2024). Lower thresholds indicate better visual acuity. As expected, thresholds were lower for unflanked stimuli (2.26’ ± 0.37’ or *≈* 20/23) and higher with flankers (2.59’ ± 0.24’ or *≈* 20/26), demonstrating a reduction in acuity in the presence of flankers (paired t-test: t(9) = −3.84, p = 0.0039, Cohen’s d = 0.96, *BF*_10_ = 13.17). This confirms that our design captures foveal crowding, consistent with previous work that used a similar stimulus design (Clark et al., 2024, Pelli et al., 2016). The magnitude of the flanker-induced threshold elevation was somewhat smaller than in prior work using a related paradigm (Clark et al., 2024), likely reflecting differences in stimulus configuration and timing, including the absence of vertical flankers and the substantially shorter stimulus duration used here. A similar pattern was observed in the pre-microsaccade condition (Figure 3*C*), where acuity thresholds remained significantly lower for unflanked stimuli (2.23’ ± 0.34’ or *≈* 20/22) than for flanked stimuli (2.87’ ± 0.39’ or *≈* 20/29), indicating robust crowding shortly before microsaccade onset. The difference between unflanked and flanked conditions was significant (paired t-test: t(9)= −10.29, p *<* 0.0001, Cohen’s d = 1.61, *BF*_10_ = 6234.81), demonstrating strong and reliable crowding effects in the pre-microsaccade interval.

**Figure 2:**
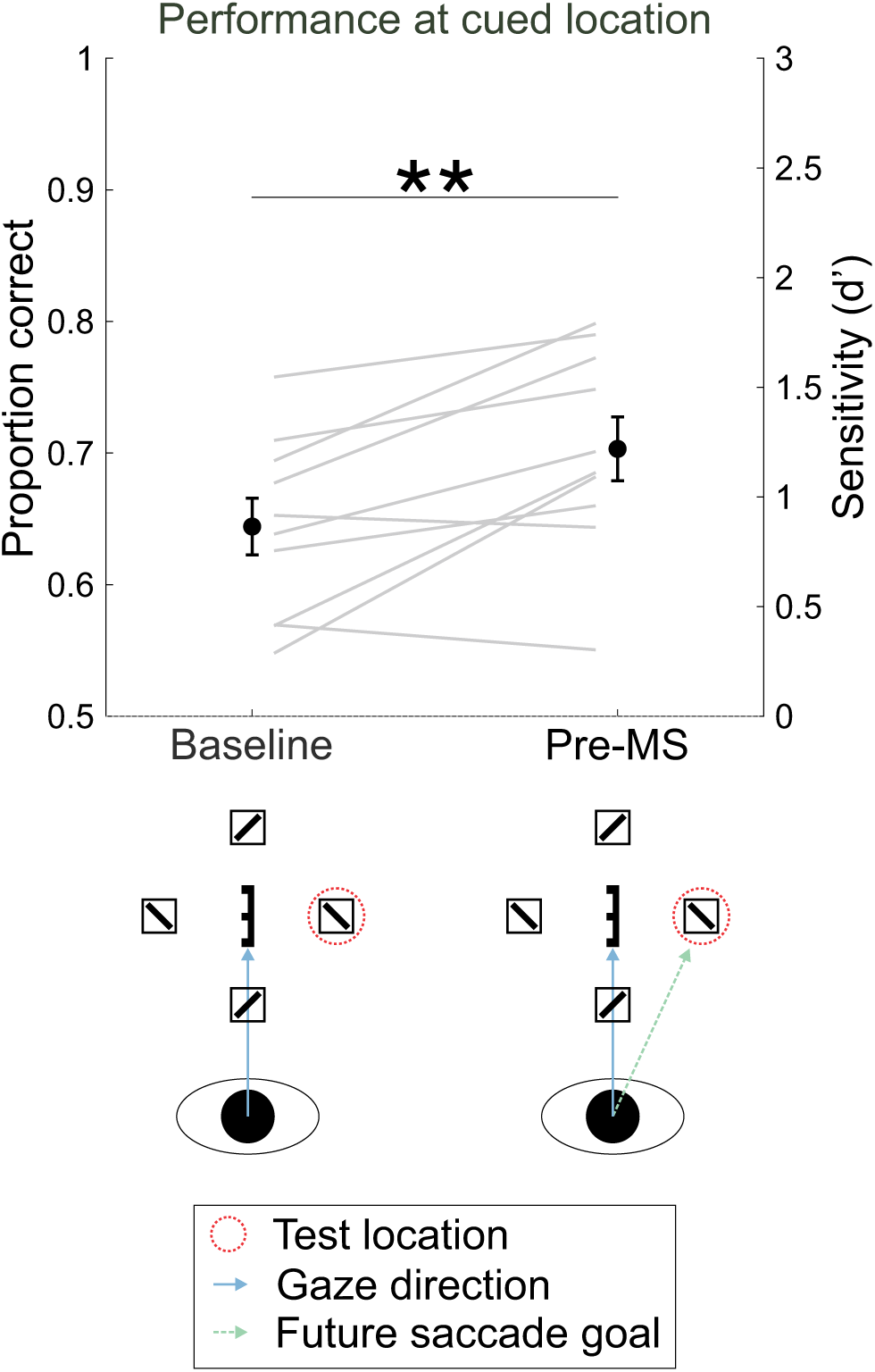
Performance at the cued location (orientation probe) across baseline and pre-microsaccade conditions. Average performance across subjects, expressed both as proportion correct (left axis) and as sensitivity index *d^′^* (right axis) for baseline and pre-microsaccade trials. Error bars represent ±1 s.e.m. Asterisks denote statistical significance between conditions (*^∗^p <* 0.05, *^∗∗^p <* 0.01, *^∗∗∗^p <* 0.001). A significant performance benefit was observed in the pre-microsaccade condition relative to baseline, consistent across both accuracy and sensitivity measures. redistribution could transiently alter the crowding zone around the PLF.

**Figure 3:**
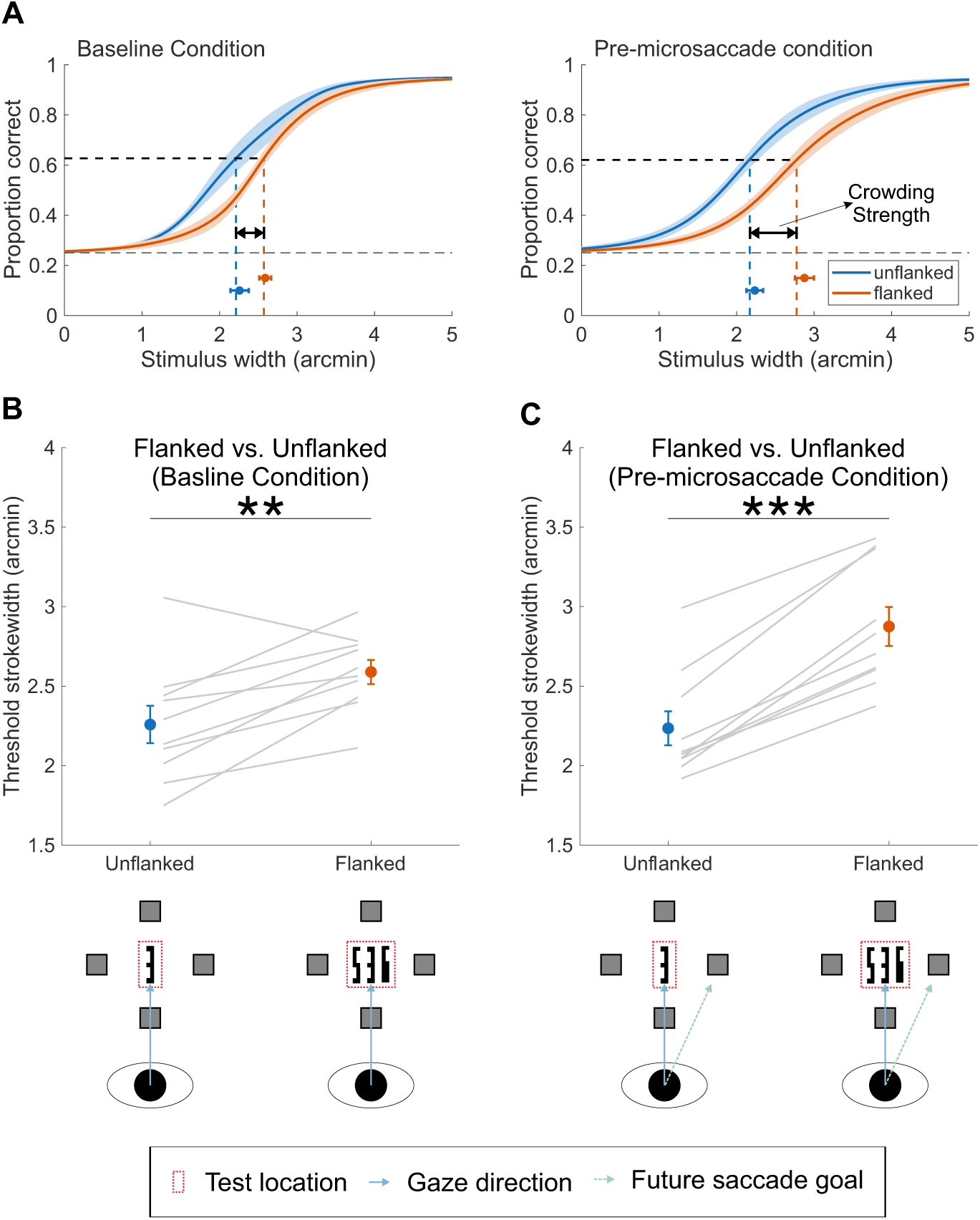
Psychometric functions and task performance for baseline and pre-microsaccade conditions. (A) Average psychometric functions across subjects showing performance in the flanked (orange) and unflanked (blue) conditions for both baseline and pre-microsaccade trials. Solid lines indicate the mean fitted curves across subjects, and the surrounding shaded areas indicating SEM. Thresholds (target stroke width at 75% correct performance) are indicated by dashed vertical lines, with the mean and SEM of these thresholds across subjects shown for each condition. (B) Target threshold stroke width for the flanked and unflanked conditions in the baseline condition. Thresholds were larger in the presence of flankers, confirming that the paradigm elicited foveal crowding. (C) Target threshold stroke width for the flanked and unflanked conditions in the pre-microsaccade condition. As in the baseline condition, thresholds were higher in the flanked configuration, demonstrating that foveal crowding persisted prior to microsaccades. Error bars represent ±1 s.e.m. Asterisks indicate statistical significance (*^∗^p <* 0.05, *^∗∗^p <* 0.01, *^∗∗∗^p <* 0.001).

To capture how much nearby flankers interfered with target identification, we quantified the flanker-induced increase in acuity threshold relative to the unflanked condition, which we refer to as crowding strength. Specifically, crowding strength was defined as ΔVA = threshold flanked – threshold unflanked, with larger values indicating stronger crowding. To this end, we compared ΔVA across the baseline and pre-microsaccade conditions (Figure 4). Crowding strength was larger in the pre-microsaccade condition (0.64*^′^ ±* 0.20*^′^*) than in the baseline condition (0.33*^′^ ±* 0.27*^′^*). Percent change was computed only for subjects with positive baseline crowding values (*n* = 9), yielding a mean 117.6 ± 50.1% increase in crowding strength. To visualize the consistency of this effect across individuals, we plotted the crowding strength for each subject in the baseline condition against the pre-microsaccade condition (Figure 4), with the dashed line indicating the unity line. Most data points fell above the unity line, indicating larger crowding strength prior to microsaccade onset. A statistical comparison of the difference relative to the unity line confirmed this increase (paired t-test on Pre-MS *−* Baseline: *t*(9) = 2.99, *p* = 0.015, Cohen’s *d* = 0.95, *BF*_10_ = 4.44). Thus, crowding strength was stronger in the pre-microsaccade condition, indicating increased crowding strength at the fovea immediately before microsaccade onset.

**Figure 4:**
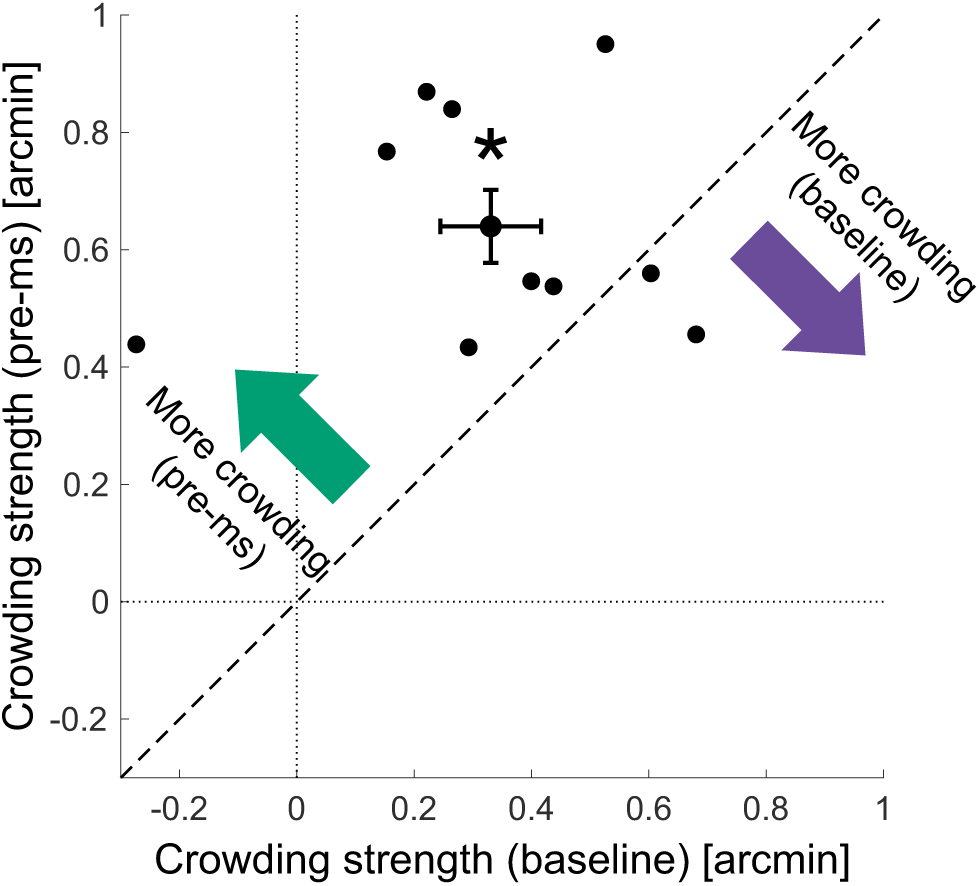
Crowding strength increases prior to microsaccade onset. Scatter plot showing crowding strength (defined as the difference in target stroke width thresholds, arcmin, between flanked and unflanked conditions) for each subject. The x-axis shows crowding strength in the baseline condition, and the y-axis shows crowding strength in the pre-microsaccade condition. Each point represents an individual subject, and the filled circle with error bars denotes the group mean ± 1 s.e.m. along both axes. The dashed line indicates the unity line. Asterisks indicate statistical significance (*^∗^p <* 0.05, *^∗∗^p <* 0.01, *^∗∗∗^p <* 0.001) of the group mean’s deviation from unity (paired t-test on pre-ms - baseline).

To test whether performance at the microsaccade goal influenced the increase in crowding observed at the PLF, we examined whether crowding strength depended on accuracy in discriminating the orientation of the stimuli presented at the microsaccade goal location (see Supplementary Figure S2). Correct responses at the cued location may reflect stronger engagement with the microsaccade goal, in which case one might expect a larger change in crowding at the PLF on correct-goal trials. We therefore fit a linear mixed-effects model with Condition (baseline vs. pre-microsaccade), goal-probe accuracy (correct vs. incorrect), and their interaction as fixed effects, and subject as a random intercept. This analysis revealed a significant main effect of Condition (*F* (1, 30) = 5.93, *p* = 0.021), confirming stronger crowding strength in the pre-microsaccade condition. However, there was no main effect of goal-probe accuracy (*F* (1, 30) = 1.06, *p* = 0.312) and no Condition *×* Accuracy interaction (*F* (1, 30) = 0.12, *p* = 0.728). The fixed-effect estimate for the pre-microsaccade condition relative to baseline was *β* = 0.31 arcmin (SE = 0.13, *t*(36) = 2.43, *p* = 0.020). Thus, crowding strength increased before microsaccade onset, but this increase did not depend on whether performance at the microsaccade goal was correct or incorrect, indicating that the effect was not explained by variation in peripheral task performance.

To quantify the spatial extent of foveal crowding, we focused on flanked trials and, for each subject and condition, estimated critical spacing as the center-to-center distance between the target and flankers at the acuity threshold (edge-to-edge spacing is a linear transformation of center-to-center spacing and yielded similar statistical outcomes). Critical spacing was significantly larger in the pre-microsaccade condition compared to baseline (paired t-test on center-to-center spacing: *t*(9) = *−*2.59, *p* = 0.029, Cohen’s *d* = 0.81, *BF*_10_ = 2.65). However, the magnitude of this difference varied across participants, with a subset of subjects exhibiting a pronounced increase in spacing thresholds prior to microsaccade onset, while others showed smaller or minimal changes. Across subjects, changes in critical spacing showed a positive rank association with changes in crowding strength (ΔVA, arcmin; Kendall’s *τ* = 0.69, *p* = 0.0047), suggesting that participants with larger pre-microsaccadic increases in acuity-based crowding also tended to show larger increases in spacing-based crowding. Together, these results provide converging evidence that the spatial extent of crowding increases prior to microsaccade onset, while also highlighting notable inter-individual variability in the spacing-based measure.

To determine whether the increased crowding strength observed in the pre-microsaccade condition reflected an expansion of pooling regions rather than increased positional uncertainty, we next examined the overall mislocalization rate (Figure 5*A*), instances where observers reported a flanker instead of the target. This was computed by combining all trials in which subjects did not identify the central target correctly, and calculating the proportion of trials where the observer reported either the left or the right flanker instead of the target. The dashed line in the figure indicates the chance level for these specific mislocalizations, corresponding to the probability of reporting one of the flankers instead of the target, given that an error was made on the location judgment in the 4AFC task (*≈* 66.7%). A paired t-test failed to detect a significant difference between conditions (t(8) = 0.54, p = 0.6041, Cohen’s dz = –0.18). The Bayesian t-test yielded a *BF*_10_ = 0.36, indicating that the data were approximately 2.8 times more likely under the null hypothesis than under the alternative hypothesis. Thus, the results provided weak evidence against the presence of an effect. This result indicates that positional uncertainty at the center of gaze remains stable between baseline and pre-microsaccade conditions, suggesting that the increase in crowding strength most likely stems from an expansion of pooling regions rather than from increased localization noise.

**Figure 5:**
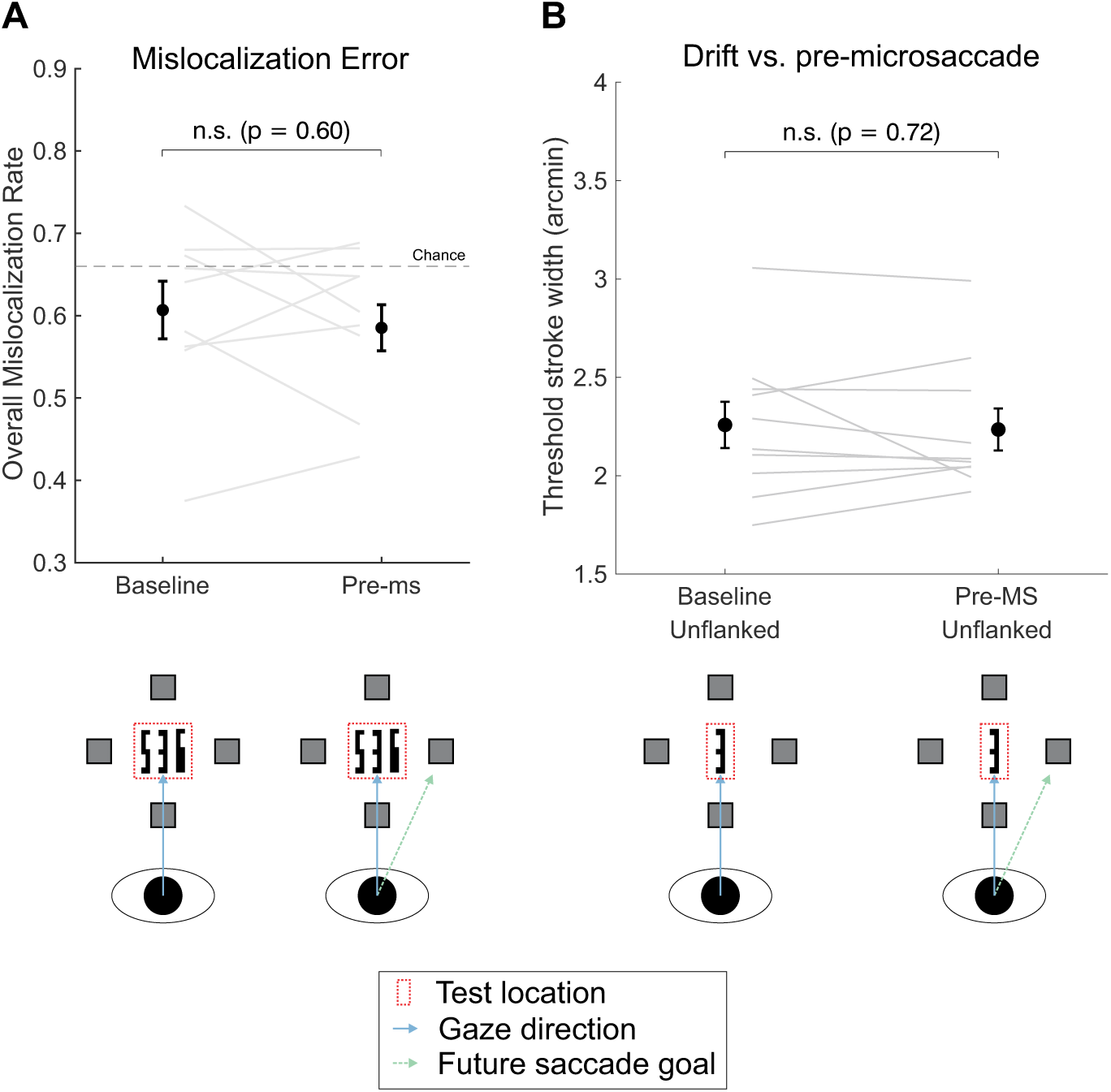
Mislocalization bias and unflanked acuity across microsaccade conditions. (A) Overall mislocalization rate (error bias) as a function of microsaccade condition (baseline vs. pre-microsaccade). The mislocalization rate reflects the combined probability of misreporting the target location as either the left or right flanker. The dashed line indicates the chance level for these specific mislocalizations, corresponding to the probability of reporting one of the flankers instead of the target, given that an error was made on the location judgment in the 4AFC task (*≈* 66.7%). (B) Change in target stroke width threshold in the unflanked condition across baseline and pre-microsaccade trials, where no significant differences were observed. Error bars represent ±1 s.e.m. Asterisks indicate statistical significance (*^∗^p <* 0.05, *^∗∗^p <* 0.01, *^∗∗∗^p <* 0.001).

Given that attentional resources are known to shift toward the future saccade goal (Guzhang et al., 2024, Hanning and Deubel, 2023, Kroell and Rolfs, 2022), we asked whether acuity at the PLF declines immediately before microsaccade onset, as reported by Guzhang et al. (2024). For this analysis, we compared unflanked acuity across the baseline and pre-microsaccade conditions, as shown in 5*B*. A paired t-test revealed no significant difference (t(9) = 0.37, p = 0.7205, Cohen’s d = 0.06, *BF*_10_ = 0.33), with comparable acuity across baseline (mean = 2.26 ± 0.37 arcmin, Snellen *≈* 20/23) and pre-microsaccade trials (mean = 2.23 ± 0.34 arcmin, Snellen *≈* 20/22). Together, these results indicate that the pre-microsaccadic increase in crowding was not accompanied by a corresponding decline in unflanked acuity at the PLF. This dissociation—stable unflanked acuity but increased crowding—indicates that microsaccade preparation can selectively alter feature integration at the PLF without producing an uniform reduction in baseline acuity.

In the current study, flankers were positioned along the radial (horizontal) axis, while subjects executed microsaccades either parallel to the flanker axis (leftward or rightward) or perpendicular to it (upward or downward). To examine whether the relative direction of the impending microsaccade influenced crowding, flanked trials from the pre-microsaccade condition were grouped by landing location, with leftward and rightward pooled as horizontal and upward and downward pooled as vertical (see Supplementary Figure S3). A paired t-test on sensitivity (d’) revealed no significant difference between horizontal and vertical microsaccade directions (t(9) = –0.75, p = 0.48, Cohen’s d = 0.25, *BF*_10_ = 0.39), indicating that crowding thresholds were not modulated by microsaccade direction.

## Discussion

Here, we investigated visual crowding at the preferred locus of fixation (PLF) while observers either maintained gaze on the crowded stimulus or were about to shift their gaze 20 arcminutes toward a cued location. Using high-precision eye tracking (Wu et al., 2023) with retinally contingent stimulus delivery (Santini et al., 2007), we ensured arcminute-level control of gaze localization during stimulus presentation and reliable detection of microsaccades. The present study provides evidence that visual crowding at the PLF is transiently exacerbated immediately prior to microsaccade onset, as reflected in a robust increase in crowding strength. This effect likely reflects a dynamic reallocation of visual processing resources during microsaccade preparation, complementing the attentional facilitation previously reported at the movement goal for spatially isolated stimuli (Guzhang et al., 2024, Shelchkova and Poletti, 2020) as well as for crowded stimuli (Prahalad and Coates, 2024).

Consistent with prior work demonstrating pre-microsaccadic perceptual benefits at the movement goal (Shelchkova and Poletti, 2020, Guzhang et al., 2024, Prahalad and Coates, 2024), the enhancement in orientation discrimination at the cued location observed here indicates that the current paradigm effectively engaged the expected pre-microsaccadic allocation of attention. The magnitude of this benefit was somewhat smaller than in some prior reports, possibly due to the requirement that observers complete the central digit report before responding to the goal stimulus, making the task more challenging. This is consistent with prior work showing that dividing attention across concurrent tasks or increasing perceptual load can reduce discrimination performance by limiting available processing resources (Pashler, 1994, Lavie, 2005). In this context, the goal-related facilitation serves primarily to establish that the paradigm successfully induced the intended shift in processing resources, providing a basis for examining how this redistribution influences visual processing at the movement start site, the PLF, which remains underexplored. More broadly, this pattern is consistent with the idea that preparing an eye movement involves a redistribution of attentional resources away from the current fixated location toward the upcoming saccade goal (Fischer and Breitmeyer, 1987).

Prior work using larger saccades has shown modulations in visual sensitivity at the PLF before movement onset. However, the nature of these changes remains unclear, with prior studies reporting contrasting findings. Some work has shown enhancements in visual performance at the fovea, particularly when stimuli are presented close to saccade onset, an effect attributed to predictive remapping of attention (Rolfs and Carrasco, 2012, Szinte et al., 2018). Kroell and Rolfs (2022) further demonstrated that such enhancements are feature-specific, occurring only for stimuli that match the saccade target, indicating that facilitation at fixation depends on task demands and stimulus properties. In contrast, Hanning and Deubel (2023) reported a drop in sensitivity at the PLF prior to movement onset using Gabor stimuli embedded in dynamic noise, with matched probes presented at both the start and goal locations. Within the foveola, Guzhang et al. (2024) provided the closest parallel, showing that fine spatial discrimination at the PLF is reduced during microsaccade preparation. However, because these measurements relied on spatially isolated probes presented sufficiently far apart to avoid crowding interactions, they do not speak to changes in feature integration or in the spatial extent of the crowding zone before the gaze shift. In light of this, a key unresolved question is whether the cost at the PLF reflects only a general reduction in sensitivity or also includes changes in how visual information is integrated across space. Our results address this question, showing that microsaccade preparation selectively increases crowding strength at the PLF without altering unflanked acuity. This pattern indicates a change in spatial integration rather than a uniform loss of sensitivity, extending previous findings by demonstrating that pre-microsaccadic modulation affects the strength and spatial extent of crowding at the start site.

Our findings are broadly consistent with those of Guzhang et al. (2024), in that both studies show reduced performance at the PLF during microsaccade preparation; however, in the current study this effect was observed specifically in the presence of flankers. However, in contrast to that study, we did not observe a change in unflanked acuity between baseline fixation and the pre-microsaccadic interval. One possible explanation for this difference may lie in the task design. First, in Guzhang et al. (2024), the response cue indicating the relevant probe was presented only after the blank interval, requiring observers to retain the orientation of all stimuli before responding, and only in a small proportion of trials were the motor and response cues congruent, such that the saccade goal did not reliably indicate the target to be reported. In contrast, in the present study, the motor and response cue were provided in advance at the movement goal location during the cueing interval (100% congruent), allowing observers to focus on the central digit and the cued orientation probe. This difference in task structure suggests that the greater overall task difficulty in the previous study may explain the stronger reduction in sensitivity at the PLF observed there. Second, Guzhang et al. (2024) used similar stimuli and task demands at both the start and goal locations, which may have increased interference across locations. In contrast, in the present study, the use of different stimuli and task demands at the two locations likely reduced such interference. Together, these differences may explain why the central sensitivity reduction was not detected in the present study.

What might underlie the increase in crowding strength at the PLF? One possibility is elevated internal noise (Lu et al., 2011, Pelli, 1990), which would be expected to reduce overall sensitivity and increase thresholds irrespective of flanker presence. However, unflanked acuity at the PLF remained stable across conditions, making a global loss of sensitivity unlikely. A second possibility is increased positional uncertainty (Greenwood et al., 2009), which would be expected to increase target–flanker mislocalizations during the pre-microsaccadic interval. We did not observe reliable differences in mislocalization across conditions, although this null result should be interpreted cautiously. Taken together, the pattern is more consistent with a change in how information is integrated across space than with a uniform degradation of visual encoding. Specifically, the increase in crowding strength is consistent with an expansion of the effective spatial integration zone at the PLF (Parkes et al., 2001, Pelli et al., 2004, Levi, 2008), leading to stronger pooling of target and flanker signals during microsaccade preparation. The increase in critical spacing, although smaller and more variable than the change in crowding strength, provides converging support for this interpretation.

Flankers were arranged along a single (horizontal) axis, with microsaccades executed either parallel or perpendicular to this configuration, providing a way to assess whether compression-related distortions during microsaccade preparation influence crowding. Peri-saccadic compression, in which visual space is transiently altered such that the perceived spatial arrangement of stimuli is compressed toward the direction of the impending saccade target, has been well documented for larger saccades and is associated with systematic distortions in the perceived locations of briefly presented stimuli just before saccade onset (Dassonville et al., 1992, Honda, 1989, Ross et al., 1997, 2001, Zirnsak et al., 2014, Weng et al., 2024). Computational accounts attribute these distortions to movement-related signals, such as efference copy from the superior colliculus, that modulate visual responses in a spatially selective manner (Hamker et al., 2008). Similar distortions have also been observed during microsaccades (Hafed, 2013). In this framework, compression would be expected to produce direction-dependent changes in crowding based on the alignment between microsaccade direction and flanker configuration. However, no such effect was observed: crowding strength was statistically comparable for microsaccades executed parallel or perpendicular to the flankers (Supplementary Figure S3). This suggests that the increase in crowding strength at the PLF is unlikely to be explained by direction-specific compression or mislocalization, and instead reflects a more global change in spatial integration during microsaccade preparation.

The interpretations from the current study are, of course, subject to certain limitations. The magnitude of the observed effect may in fact represent an underestimate, as observers consistently expected a crowded stimulus at the PLF, potentially biasing processing toward the center. While this approach was adopted to maximize the number of valid trials, introducing occasional catch trials in which the central stimulus is omitted could further reduce the need to divide processing resources between locations and may reveal a stronger effect. Despite this, we observed a robust increase in crowding strength within the pre-microsaccadic interval examined here, suggesting that the true magnitude of modulation at the PLF may be even larger.

In conclusion, microsaccade preparation increases visual crowding at the PLF while leaving unflanked acuity unchanged. This finding shows that the perceptual consequences of microsaccade preparation are not limited to facilitation at the upcoming gaze target (Prahalad and Coates, 2024, Guzhang et al., 2024, Shelchkova and Poletti, 2020). Before the eyes move, spatial integration is also altered at the current fixation location, reducing the ability to segregate nearby objects at the center of gaze. Because microsaccades occur frequently during high-acuity tasks such as reading, face recognition, and fine spatial discrimination (Poletti et al., 2013, Intoy and Rucci, 2020, Ko et al., 2010, Bowers and Poletti, 2017, Shelchkova et al., 2019, Clark et al., 2022), these results reveal a dynamic reorganization of foveal processing during fixation itself. More broadly, they show that microsaccade preparation modulates not only visual sensitivity, but also the spatial integration mechanisms that determine object segregation in the foveola.

## Methods

### Observers

Twelve adult observers (*≥*18 years; age range: 20–30 years) with normal or corrected-to-normal vision (20/20 or better) participated in the study. The cohort comprised nine females and three males. No participants withdrew after enrollment. Two participants were excluded from further analysis due to unreliable psychometric fits and unstable threshold estimates that did not meet predefined data-quality criteria (see Data Analysis). The final dataset therefore included ten observers: five näıve individuals and five experienced observers who had previously participated in unrelated eyetracking experiments, one of whom was an author; except for the author, all observers were näıve to the purpose of the study. An a priori power analysis was conducted using G*Power (Kang, 2021) to determine the required sample size for detecting changes in crowding strength between baseline and pre-microsaccadic conditions. Effect size estimates were informed by prior studies reporting robust pre-microsaccadic modulations of fine spatial discrimination and crowding-related interference at the foveal level, which have consistently shown large within-subject effects (e.g., Prahalad et al., 2025, Guzhang et al., 2024). Assuming a large effect size (*d_z_ ≈* 0.8–1.0), a minimum sample size of 9–15 observers was estimated to achieve 80% power for a paired-samples comparison (*α* = 0.05, two-tailed). The experiment was performed monocularly and the non-tested eye was occluded with an eye patch. An initial screening was performed, all participants demonstrated visual acuity of 20/20 or better with or without the need for corrective lenses. The experiment protocol was approved by the University of Rochester’s Research subjects review board (RSRB). Written informed consent was obtained from each participant after explaining the study procedures and reviewing the consent form.

### Stimuli and Apparatus

Stimuli were displayed on an LCD monitor (ASUS PG259QN), with a vertical refresh rate of 360 Hz and a spatial resolution of 1920 *×* 1080. The monitor was positioned at *≈* 5 meters from the observer where each pixel subtended *≈* 0.19 arcmin. Eye movements were recorded with high precision using a custom-digital Dual Purkinje Image (dDPI) eye tracker (Wu et al., 2023), with a sampling rate of 1000 Hz. The observer’s head was immobilized with a dental-imprint bite bar and a magnetized helmet attached to a head holder, which helped minimize the noise from head movement and enhanced eye-tracking precision.

The central digit stimulus consisted of digits 3, 5, 6 and 9 from the Pelli number font (Pelli et al., 2016), a font specifically designed to test visual crowding at the fovea. It has a vertically elongated design (aspect ratio 5:1), which allows for testing at smaller center to center spacing between the target and the flanker when compared to traditional optotypes with equal aspect ratio. Stimuli were presented at maximum contrast in black text on an uniform gray background and were controlled using EyeRIS (Santini et al., 2007), a custom-developed system enabling gaze-contingent display control. Stimuli were presented either in isolation (unflanked) or were presented alongside flankers (flanked) which were positioned along the radial axis (horizontal). In flanked trials, target size was staircase-controlled and flanker spacing was maintained at a fixed spacing-to-size ratio (1.4×), so critical spacing was computed as the center-to-center distance at threshold.

Observers were instructed to fixate centrally, surrounded by four square placeholders (7.5*^′^ ×* 7.5*^′^*) positioned in a circle with a radius of 20 arcminutes at the cardinal positions (see figure 1*A*). Additionally, a line orientation probe ( ±45*^◦^*) was presented simultaneously within each square placeholder as a secondary stimulus. Because the Pelli digits are vertically elongated, flankers positioned along the tangential (vertical) axis would be farther from the target, potentially overlapping with the placeholders and the orientation probe, thereby limiting the range of flanker spacings that could be tested. Thus, flankers were positioned only along the radial (horizontal) axis. There were two experimental conditions in this study: (1) baseline condition, where observers were instructed to maintain fixation at the center of the monitor throughout the trial, and (2) pre-microsaccade condition, where observers were instructed to shift their gaze to the square placeholder that changed color. Flanked and unflanked trials were randomly interleaved within each block, whereas the baseline and pre-microsaccade conditions were presented in separate, dedicated blocks. In the first session, all observers completed a fixed training sequence (baseline/drift block followed by a pre-microsaccade block) to familiarize them with task demands and the cueing paradigm. In subsequent sessions, block order (baseline vs. pre-microsaccade) was randomized across observers.

### Experimental Paradigm

Data collection involved multiple experimental sessions, with an average of 5 sessions per subject. Each session lasted approximately one hour, with 5-6 blocks of data collected per session. At the beginning of each block, a two-step gaze-contingent calibration was performed to accurately localize the observer’s gaze direction. The calibration process included two phases: During the first phase (automatic calibration), observers sequentially fixated on a 3×3 grid of points. In the second phase (manual calibration), observers refined the calibration by fixating on the same set of points while manually adjusting a gaze-contingent marker to overlap with each point. The manual calibration step was repeated only for the central point of the grid at the beginning of each trial to account for potential head movement between trials.

Each trial began with a fixation period (randomly chosen between 500 - 800 ms), during which the five square placeholders were presented. Observers were instructed to maintain fixation on the central square placeholder. After the fixation interval, the central square placeholder disappeared, and one of the four square placeholders at the cardinal positions changed its line color from black to red, serving as a motor cue. The stimulus was presented for 100 ms and consisted of the crowded stimulus (in Pelli font) and the high-acuity tilted lines presented within each square placeholder. Following the stimulus interval, a blank interval of 500 ms was introduced, during which only the square placeholders remained on the screen across both conditions. This allowed observers to make the gaze shift in the pre-microsaccade condition, while they maintained fixation on the central square placeholder in the baseline condition. After the blank interval, observers were instructed to report the number displayed in the center of the crowded stimulus. Following this, the square placeholders were displayed again, and observers were instructed to report the orientation of the tilted line, but only for the one in the red box. Subjects were always required to report the orientation of the tilted line within the cued box (100 % validity). The experiment sequence is illustrated in figure 1*A*.

Target acuity and crowding thresholds (critical spacing) were determined using the Parametric Estimation by Sequential Testing (PEST) procedure (Taylor and Creelman, 1967), in which target size was adaptively adjusted based on the observer’s performance to converge at 62.5% correct in a 4-AFC task. In flanked trials, center-to-center flanker spacing was fixed at 1.4 times the target size, such that the absolute spacing varied as target size was modulated during the staircase. In the pre-microsaccade condition, we aimed to present the stimulus prior to the onset of the microsaccade to increase the efficiency of the experiment. To increase the efficiency of the experimental paradigm and maximize the number of trials with stimuli presented prior to microsaccade onset in the pre-microsaccade condition, we adopted methods similar to previous studies (Prahalad and Coates, 2024, Hunt and Cavanagh, 2011). Microsaccade latency was estimated at the end of each block by detecting microsaccades that occurred following the motor cue onset. Microsaccades were identified using an angular-velocity threshold of 4.5 arcmin/frame (minimum duration *≥* 3 samples *≈* 8 ms), with events classified only if no saccades occurred between cue and stimulus onset. This allowed us to compute the median saccadic latency (SRT) based on real-time onset estimates. The cue-to-stimulus onset time was then determined using the median microsaccade latency, and for each trial, we manipulated the stimulus onset time by subtracting 25, 50, or 75 ms from the SRT.

### Refractive Correction

We employed a Badal lens setup to compensate for spherical refractive error among observers. This system was used for all observers, regardless of whether they required refractive correction, to ensure consistency across participants. To guide the correction process, we used Atchison’s model (Atchison et al., 1995), which describes a relay system that achieves accurate correction with minimal distortion.

To assess visual acuity, we presented a duo-chrome test using a tumbling E chart with 20/15, 20/20 and 20/32 lines. Participants adjusted the distance between the lenses until the target appeared equally clear on both the red and green backgrounds. Visual acuity was measured at this endpoint, once magnification was accounted for, and was ensured to be 20/20 or better. This approach ensured precise correction for all participants, maintaining high visual acuity.

### Data Analysis

Eye movements were automatically classified as saccades and drifts, with all classifications further validated manually by trained laboratory personnel. Investigators were not blinded to experimental condition during data collection. Trials were discarded if any blink events occurred during the trial, if eye position during stimulus presentation drifted more than 15 arcminutes from the center of the display (with a tolerance of up to 5 samples outside this range), or if saccadic eye movements occurred between cue onset and stimulus onset. In the baseline condition, trials further required no microsaccadic events for at least 300 ms following stimulus onset.

Figure 1*B* shows an example eye trace from a pre-microsaccadic trial, while Figure 1*C* shows the distribution of microsaccade onset times relative to stimulus onset. Pre-microsaccade trials were defined as those in which the microsaccade occurred 0–350 ms after stimulus onset. These trials were then further filtered using the following additional criteria: (1) a valid microsaccade was required to occur within the designated time window following stimulus onset; (2) after the selected microsaccade ended, the mean gaze position in the 10–30 ms interval following microsaccade offset was required to fall within 15 arcmin from the center of the cued location; and (3) trials were discarded if another microsaccade occurred within 100 ms of the selected one. Finally, Figure 1*D* shows the amplitude distribution of microsaccades for the valid pre-microsaccade trials that passed these criteria.

Visual acuity was assessed both in terms of stimulus width (in arcminutes) and the minimum angle of resolution (MAR). The visual acuity threshold, defined as the smallest stimulus width for reliable performance above chance (62.5% correct, with a 25% chance level), was calculated using a logistic function implemented in psignifit (v4.1.10; Wichmann Lab GitHub repository; Schütt et al. 2016). A lower threshold indicates better acuity. To convert the Pelli digits stroke width to MAR, instead of using 1/5 of the stimulus width as with the tumbling E, MAR was defined as 1/2 of the stimulus stroke width (Pelli et al., 2016). Thus, an optotype with a width of 2 arcmin would correspond to the 20/20 Snellen MAR line.

To evaluate the reliability of psychometric fits, we computed multiple goodness-of-fit metrics (deviance, log-likelihood, AIC, and BIC) for each subject. Individual values were compared against the group distribution, and subjects whose fits fell more than 2 standard deviations from the group mean on any of these metrics were classified as outliers. Using this procedure, 2 of the 12 observers were excluded, yielding a final sample of 10 observers for the main analyses.

### Statistical Analysis

Statistical analyses were performed in MATLAB (MathWorks, Natick, MA; RRID:SCR 001622) using the Statistics and Machine Learning Toolbox. Planned pairwise comparisons between conditions were conducted using paired two-tailed t-tests. To assess the robustness of these comparisons to distributional assumptions, we additionally performed non-parametric Wilcoxon signed-rank tests, which yielded results consistent with the corresponding parametric tests. Each comparison was accompanied by the calculation of Bayes factors (*BF*_10_; Bayes Factor Matlab toolbox (GitHub repository)) and Cohen’s *D* to quantify the strength and magnitude of the observed effects. Repeated-measures analyses of variance (RANOVA) were used for factorial comparisons, with all factors treated as within-subject variables. For repeated-measures factors with more than two levels, Mauchly’s test was used to assess violations of the sphericity assumption; when violated, Greenhouse-Geisser-corrected p-values are reported. For two-level within-subject factors, sphericity corrections were unnecessary.

To complement paired comparisons, we also analyzed the data using linear mixed-effects models (LME) with condition as a fixed effect and subject as a random intercept. This approach accounts for within-subject dependence and individual variability while providing population-level estimates of condition effects. Models were fit using restricted maximum likelihood, and inference on fixed effects was based on Satterthwaite-approximated degrees of freedom. The overall findings remained consistent between the analysis methods (see supplementary table 2).

## RESOURCE AVAILABILITY

### Lead contact

Further information and requests for resources should be directed to and will be fulfilled by the lead contact, Martina Poletti.

### Data and code availability

- All original behavioral data supporting the findings of this study are publicly available via the Open Science Framework (OSF; DOI:10.17605/OSF.IO/RFZ2E).
- All custom code used for data analysis and figure generation is publicly available via the same OSF repository (DOI:10.17605/OSF.IO/RFZ2E).
- Any additional information required to reanalyze the data reported in this paper is available from the lead contact upon reasonable request.

## Acknowledgments

This work was supported by the National Institutes of Health grants R01EY029788-01 (to M.P.) and by NIH EY001319 grant to the Center for Visual Science at the University of Rochester. The authors thank Zoe Stearns for helpful comments on an earlier version of the manuscript.

## Author Contributions

K.S.P. and M.P. designed the study. K.S.P. collected the data. K.S.P. and M.P. analyzed the data and wrote the manuscript.

## Supplementary Materials

**Table 1:** Summary of valid and rejected trials. Number of valid trials and trials rejected for each criterion in the baseline and pre-microsaccade conditions. Rejection categories are reported separately and may overlap.

| Subject | Condition | Valid Trials | Total Trials | MS during stimulus presentation | MS between cue and stimulus onset | Exceeded gaze limits |
| --- | --- | --- | --- | --- | --- | --- |
| S1 | Baseline | 510 | 577 | 34 | 34 | 2 |
| S2 | Baseline | 587 | 676 | 49 | 39 | 5 |
| S3 | Baseline | 346 | 366 | 7 | 10 | 4 |
| S4 | Baseline | 405 | 566 | 99 | 69 | 11 |
| S5 | Baseline | 206 | 411 | 14 | 9 | 183 |
| S6 | Baseline | 257 | 294 | 9 | 10 | 20 |
| S7 | Baseline | 121 | 152 | 9 | 19 | 6 |
| S8 | Baseline | 361 | 377 | 10 | 6 | 0 |
| S9 | Baseline | 340 | 410 | 16 | 49 | 8 |
| S10 | Baseline | 295 | 390 | 38 | 69 | 6 |
| S1 | Pre-MS | 250 | 700 | 335 | 67 | 70 |
| S2 | Pre-MS | 270 | 592 | 252 | 87 | 57 |
| S3 | Pre-MS | 510 | 1561 | 572 | 417 | 364 |
| S4 | Pre-MS | 207 | 953 | 421 | 270 | 70 |
| S5 | Pre-MS | 191 | 884 | 465 | 45 | 199 |
| S6 | Pre-MS | 162 | 1061 | 728 | 119 | 135 |
| S7 | Pre-MS | 174 | 1054 | 829 | 263 | 56 |
| S8 | Pre-MS | 453 | 1014 | 508 | 42 | 39 |
| S9 | Pre-MS | 339 | 740 | 260 | 100 | 64 |
| S10 | Pre-MS | 249 | 665 | 287 | 93 | 38 |

**Figure S1:**
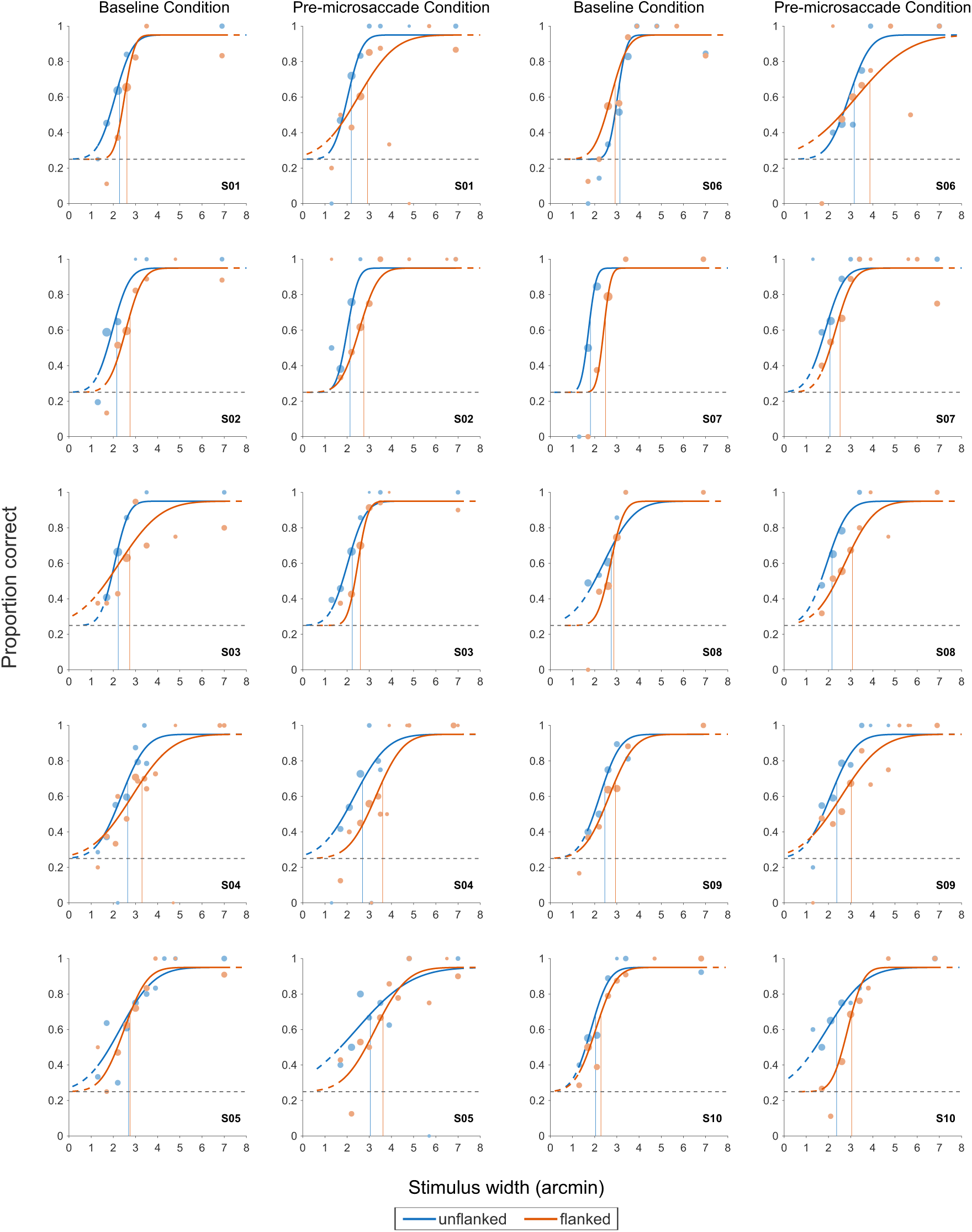
Individual psychometric functions for baseline and pre-microsaccade conditions. Psychometric functions are shown separately for each subject in the baseline and pre-microsaccade conditions. Blue symbols and curves indicate the unflanked condition, and orange symbols and curves indicate the flanked condition. Each panel shows the fitted psychometric functions relating target stroke width to proportion correct, with vertical lines indicating the estimated threshold stroke width for each condition. The horizontal black line indicates the 4AFC guess rate of 25%.

**Figure S2:**
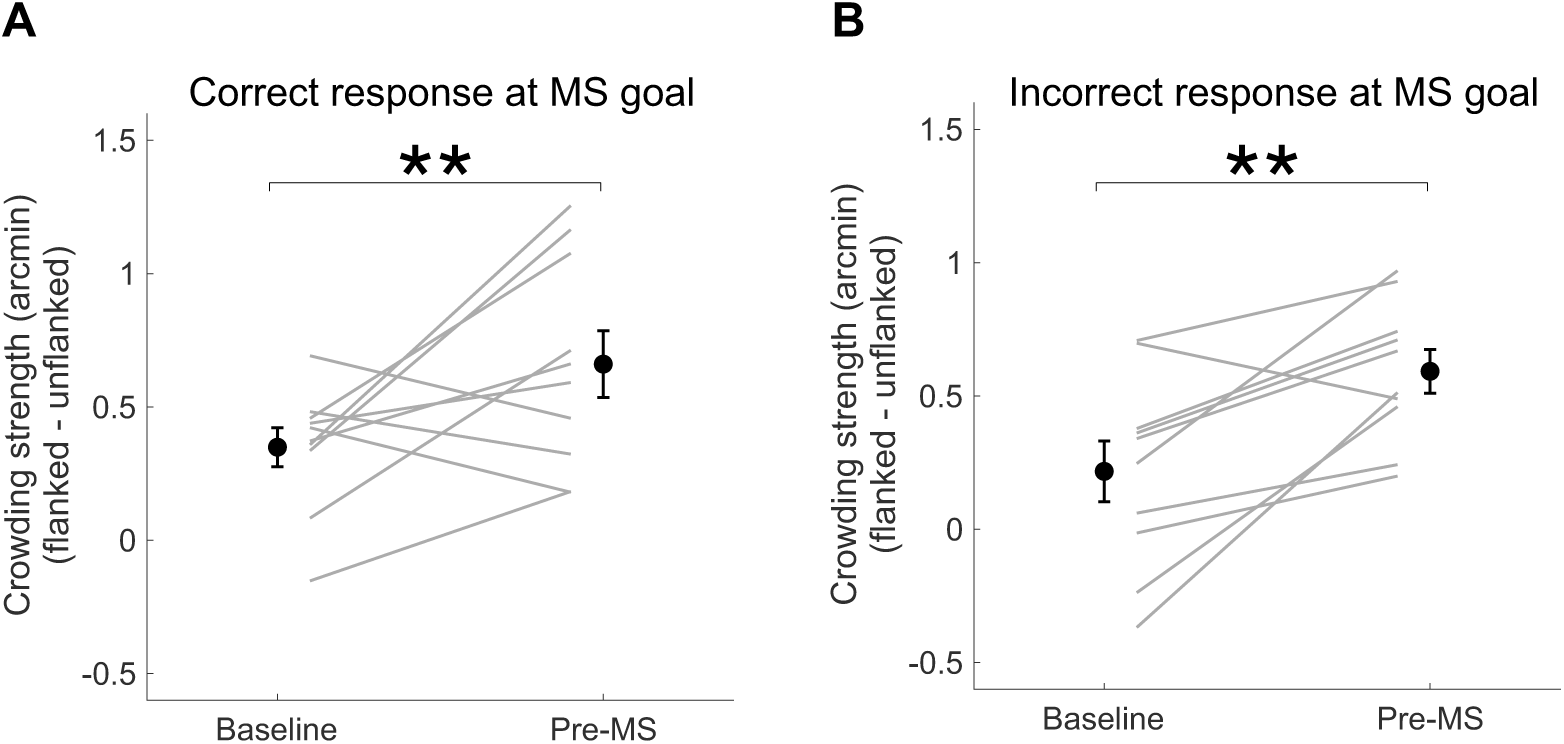
Crowding strength based on accuracy at the microsaccade goal. (A) Difference in target stroke width thresholds (in arcmin) between the flanked and unflanked conditions, computed from the subset of trials in which subjects responded *correctly* to the stimulus presented at the microsaccade goal. (B) Same measure computed from trials in which subjects responded *incorrectly* to the goal stimulus. Error bars represent ±1 s.e.m. Asterisks indicate statistical significance (*^∗^p <* 0.05, *^∗∗^p <* 0.01, *^∗∗∗^p <* 0.001).

**Figure S3:**
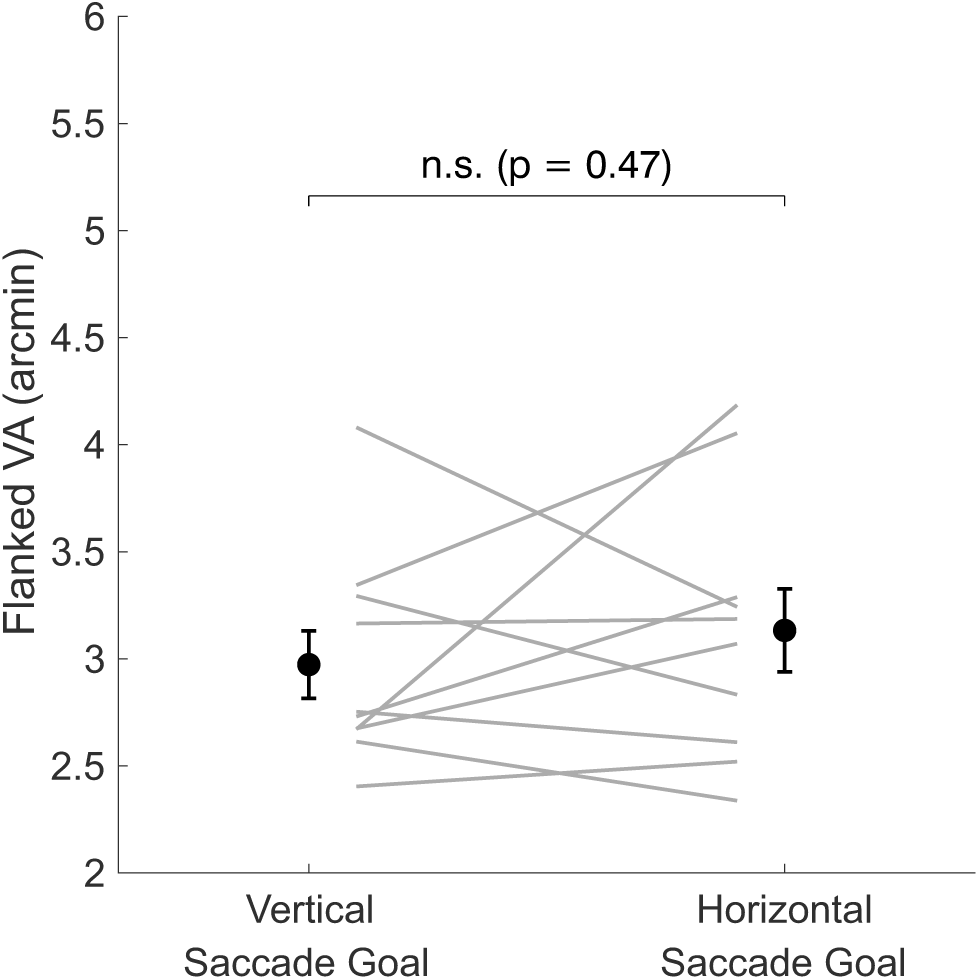
Crowding thresholds by microsaccade direction. Thresholds in arcmin for flanked trials as a function of microsaccade direction relative to the flanker configuration. Thresholds are shown separately for microsaccades parallel to the flanker axis (horizontal) and perpendicular to the flanker axis (vertical). Error bars represent ±1 s.e.m. Asterisks indicate statistical significance (*^∗^p <* 0.05,*^∗∗^p <* 0.01, *^∗∗∗^p <* 0.001).

**Table 2:** Summary of linear mixed-effects (LME) models. Fixed-effect estimates for the main comparisons reported in the study. All models included Condition as a fixed effect and Subject as a random intercept.

| Figure | Outcome | $\beta$ | SE | $t$ | $p$ | 95% CI |
| --- | --- | --- | --- | --- | --- | --- |
| Fig. 2 | $d'$ (goal performance) | 0.35 | 0.09 | 3.75 | 0.0015 | [0.16, 0.55] |
| Fig. 3B | Threshold (arcmin) | 0.331 | 0.082 | 4.05 | 0.0007 | [0.159, 0.502] |
| Fig. 3C | Threshold (arcmin) | 0.640 | 0.059 | 10.85 | < 0.0001 | [0.516, 0.764] |
| Fig. 4 | Pre-MS vs Drift VA change | 0.3092 | 0.0981 | 3.152 | 0.0055 | [0.1031, 0.5153] |
| Fig. 5A | Mislocalization rate | -0.014 | 0.025 | -0.57 | 0.575 | [-0.0676, 0.0388] |
| Fig. 5B | Unflanked threshold (arcmin) | -0.023 | 0.060 | -0.39 | 0.702 | [-0.150, 0.10] |

## References

Bowers, N. R. and Poletti, M. (2017). Microsaccades during reading. PLoS ONE, 12(9):e0185180.

Ko, H.-k., Poletti, M., and Rucci, M. (2010). Microsaccades precisely relocate gaze in a high visual acuity task. Nat. Neurosci., 13(12):1549–1553.

Shelchkova, N., Tang, C., and Poletti, M. (2019). Task-driven visual exploration at the foveal scale. Proc. Natl. Acad. Sci. USA, 116(12):5811–5818.

Intoy, J. and Rucci, M. (2020). Finely tuned eye movements enhance visual acuity. Nat. Commun., 11(1):795.

Clark, A. M., Intoy, J., Rucci, M., and Poletti, M. (2022). Eye drift during fixation predicts visual acuity. Proc. Natl. Acad. Sci. USA, 119(49):e2200256119.

Rucci, M. and Poletti, M. (2015). Control and functions of fixational eye movements. Annu. Rev. Vis. Sci., 1:499–518.

Guzhang, Y., Shelchkova, N., Clark, A. M., and Poletti, M. (2024). Ultra-fine resolution of pre-saccadic attention in the fovea. Curr. Biol., 34(2):309–321.e4.

Shelchkova, N. and Poletti, M. (2020). Modulations of foveal vision associated with microsaccade preparation. Proc. Natl. Acad. Sci. USA, page 201919832.

Prahalad, K. S. and Coates, D. R. (2024). Alterations to foveal crowding with microsaccade preparation. Vision Res., 214:108338.

Hanning, N. M. and Deubel, H. (2023). A dynamic 1/f noise protocol to assess visual attention without biasing perceptual processing. Behav. Res. Methods, 55(3):1503–1518.

Kroell, L. and Rolfs, M. (2022). Foveal vision anticipates defining features of eye movement targets. eLife, 11.

Stuart, J. A. and Burian, H. M. (1962). A Study of Separation Difficulty: Its Relationship to Visual Acuity in Normal and Amblyopic Eyes. Am. J. Ophthalmol., 53(3):471–477.

Bouma, H. (1970). Interaction effects in parafoveal letter recognition. Nature, 226(5241):177–178.

Levi, D. M. (2008). Crowding—An essential bottleneck for object recognition: A mini-review. Vision Res., 48(5):635–654.

Pelli, D. G. (2008). Crowding: A cortical constraint on object recognition. Curr. Opin. Neurobiol., 18(4):445–451.

Whitney, D. and Levi, D. M. (2011). Visual crowding: a fundamental limit on conscious perception and object recognition. Trends Cogn. Sci., 15(4):160–168.

Flom, M. C., Weymouth, F. W., and Kahneman, D. (1963). Visual Resolution and Contour Interaction. JOSA, 53(9):1026–1032.

Coates, D. R. and Levi, D. M. (2014). Contour interaction in foveal vision: a response to siderov, waugh, and bedell (2013). Vision Res., 96:140–144.

Siderov, J., Waugh, S. J., and Bedell, H. E. (2013). Foveal contour interaction for low contrast acuity targets. Vision Res., 77:10–13.

Bondarko, V. M., Chikhman, V. N., Danilova, M. V., and Solnushkin, S. D. (2024). Foveal crowding for large and small Landolt Cs: Similarity and Attention. Vision Res., 215:108346.

Clark, A. M., Huynh, A., and Poletti, M. (2024). Oculomotor Contributions to Foveal Crowding. J. Neurosci., 44(48).

Shamsi, F., Liu, R., and Kwon, M. (2022). Foveal crowding appears to be robust to normal aging and glaucoma unlike parafoveal and peripheral crowding. J. Vis., 22(8):10.

Prahalad, K. S., Clark, A. M., Moon, B., Roorda, A., Tiruveedhula, P., Harmening, W., Gutnikov, A., Jenks, S. K., Kapisthalam, S., Rucci, M., Rolland, J. P., and Poletti, M. (2025). Non-uniform signal pooling across the foveola. Curr. Biol., 35(6):1093–1105.

Manassi, M., Sayim, B., and Herzog, M. H. (2012). Grouping, pooling, and when bigger is better in visual crowding. J. Vis., 12(10):13.

Manassi, M., Sayim, B., and Herzog, M. H. (2013). When crowding of crowding leads to uncrowding. J. Vis., 13(13):10.

Herzog, M. H., Sayim, B., Chicherov, V., and Manassi, M. (2015). Crowding, grouping, and object recognition: A matter of appearance. J. Vis., 15(6):5.

Pelli, D. G., Waugh, S. J., Martelli, M., Crutch, S. J., Primativo, S., Yong, K. X., Rhodes, M., Yee, K., Wu, X., Famira, H. F., and Yiltiz, H. (2016). A clinical test for visual crowding. F1000Res., 5:81.

Harrison, W. J., Mattingley, J. B., and Remington, R. W. (2013). Eye Movement Targets Are Released from Visual Crowding. J. Neurosci., 33(7):2927–2933.

Lin, H., Rizak, J. D., Ma, Y.-y., Yang, S.-c., Chen, L., and Hu, X.-t. (2014). Face Recognition Increases during Saccade Preparation. PLOS ONE, 9(3):e93112.

Wolfe, B. A. and Whitney, D. (2014). Facilitating recognition of crowded faces with presaccadic attention. Front. Hum. Neurosci., 8.

Ăgaŏglu, M. N., Öğmen, H., and Chung, S. T. (2016). Unmasking saccadic uncrowding. Vision Res., 127:152–164.

Ăgaŏglu, M. N. and Chung, S. T. (2017). Interaction between stimulus contrast and pre-saccadic crowding. R. Soc. Open Sci., 4(2):160559.

Buonocore, A., Fracasso, A., and Melcher, D. (2017). Pre-saccadic perception: Separate time courses for enhancement and spatial pooling at the saccade target. PLOS ONE, 12(6):e0178902.

Lu, Z.-L., Hua, T., Huang, C.-B., Zhou, Y., and Dosher, B. A. (2011). Perceptual learning reflects external noise filtering and internal noise reduction through channel reweighting. Proc. Natl. Acad. Sci. USA, 108(25):10128–10133.

Pelli, D. G. (1990). The quantum efficiency of vision. Cambring University Press, 30(11):1811–1820.

Greenwood, J. A., Bex, P. J., and Dakin, S. C. (2009). Positional averaging explains crowding with letter-like stimuli. Proc. Natl. Acad. Sci. USA, 106(31):13130–13135.

Parkes, L., Lund, J., Angelucci, A., Solomon, J. A., and Morgan, M. (2001). Compulsory averaging of crowded orientation signals in human vision. Nat. Neurosci., 4(7):739–744.

Pelli, D. G., Palomares, M., and Majaj, N. J. (2004). Crowding is unlike ordinary masking: Distinguishing feature integration from detection. J. Vis., 4(12):12.

Wu, R.-J., Clark, A. M., Cox, M. A., Intoy, J., Jolly, P. C., Zhao, Z., and Rucci, M. (2023). High-resolution eye-tracking via digital imaging of Purkinje reflections. J. Vis., 23(5):4.

Santini, F., Redner, G., Iovin, R., and Rucci, M. (2007). EyeRIS: A general-purpose system for eye-movement-contingent display control. Behav. Res. Methods, 39(3):350–364.

Poletti, M. and Rucci, M. (2016). A compact field guide to the study of microsaccades: Challenges and functions. Vision Res., 118:83–97.

Pashler, H. (1994). Dual-task interference in simple tasks: Data and theory. Quarterly Journal of Experimental Psychology Section A, 47(2):221–256.

Lavie, N. (2005). Distracted and confused?: Selective attention under load. Trends in Cognitive Sciences, 9(2):75–82.

Fischer, B. and Breitmeyer, B. (1987). Mechanisms of visual attention revealed by saccadic eye movements. Neuropsychologia, 25(1):73–83.

Rolfs, M. and Carrasco, M. (2012). Rapid simultaneous enhancement of visual sensitivity and perceived contrast during saccade preparation. J. Neurosci., 32(40):13744–13752a.

Szinte, M., Jonikaitis, D., Rangelov, D., and Deubel, H. (2018). Pre-saccadic remapping relies on dynamics of spatial attention. Elife, 7:e37598.

Dassonville, P., Schlag, J., and Schlag-Rey, M. (1992). Oculomotor localization relies on a damped representation of saccadic eye displacement in human and nonhuman primates. Visual Neuroscience, 9(3-4):261–269.

Honda, H. (1989). Perceptual localization of visual stimuli flashed during saccades. Percept. Psychophys., 45(2):162–174.

Ross, J., Morrone, M. C., and Burr, D. C. (1997). Compression of visual space before saccades. Nature, 386(6625):598–601.

Ross, J., Morrone, M., Goldberg, M. E., and Burr, D. C. (2001). Changes in visual perception at the time of saccades. Trends Neurosci., 24(2):113–121.

Zirnsak, M., Steinmetz, N. A., Noudoost, B., Xu, K. Z., and Moore, T. (2014). Visual space is compressed in prefrontal cortex before eye movements. Nature, 507(7493):504–507.

Weng, G., Akbarian, A., Clark, K., Noudoost, B., and Nategh, N. (2024). Neural correlates of perisaccadic visual mislocalization in extrastriate cortex. Nature Communications, 15(1):6335.

Hamker, F. H., Zirnsak, M., Calow, D., and Lappe, M. (2008). The Peri-Saccadic Perception of Objects and Space. PLoS Computational Biology, 4(2).

Hafed, Z. (2013). Alteration of Visual Perception prior to Microsaccades. Neuron, 77(4):775–786.

Poletti, M., Listorti, C., and Rucci, M. (2013). Microscopic Eye Movements Compensate for Nonho-mogeneous Vision within the Fovea. Curr. Biol., 23(17):1691–1695.

Kang, H. (2021). Sample size determination and power analysis using the g*power software. J. Educ. Eval. Health Prof., 18:17.

Taylor, M. M. and Creelman, C. D. (1967). PEST: Efficient Estimates on Probability Functions. J. Acoust. Soc. Am., 41(4A):782–787.

Hunt, A. R. and Cavanagh, P. (2011). Remapped visual masking. J. Vis., 11(1):13–13.

Atchison, D. A., Bradley, A., Thibos, L. N., and Smith, G. (1995). Useful variations of the Badal optometer. Optom. Vis. Sci., 72(4):279–284.

Schütt, H. H., Harmeling, S., Macke, J. H., and Wichmann, F. A. (2016). Painfree and accurate Bayesian estimation of psychometric functions for (potentially) overdispersed data. Vision Res., 122:105–123.

